## Supplementary materials for "Epigenetic landscape of pancreatic neuroendocrine tumours reveals distinct cells of origin and means of tumour progression"

**Supplementary tables**

**Table S1:** Clinico-pathological and molecular characteristics of PanNET cohort 1

**Table S2:** Differentially methylated CpG sites identified comparing normal $\alpha$- to normal $\beta$-cells

**Table S3:** Clinico-pathological and molecular characteristics of cohort 2

**Table S4:** Differentially methylated CpG sites identified comparing $\alpha$-like to $\beta$-like PanNETs

**Table S5:** Differentially methylated CpG sites identified comparing $\alpha$-like to intermediate PanNETs

**Table S6:** Differentially methylated CpG sites identified comparing intermediate to $\beta$-like PanNETs

**Table S7:** Clinico-pathological and molecular characteristics of Chan *et al.* cohort

#### Supplementary figure legends

**Figure S1.** Flow-chart indicating number of patients used for each analysis.

**Figure S2. A.** Phyloepigenetic analysis of PanNET and normal α- and β-cell samples. Rooted tree was created with an arbitrary chosen α-sample as the root and according to the differentially methylated CpGs between sorted normal α- and β-cell samples (n=2131, adj. p-value<0.001 and $|\Delta\beta|>0.2$). In the blue and orange squares are included the α-like and β-like tumours, respectively. **B.** Cell type contributions of sorted exocrine (acinar, duct, pancreatic fibroblasts) and endocrine (α- and β-cells) pancreatic cells and blood cells (granulocytes, CD4+ and CD8+ T cells, CD14+ Monocytes, CD19+ B cells, CD56+ natural killer cells, neutrophils and eosinophils cells), in each PanNET sample based on methylation profiles. Scale refers to the percentage of contribution of each sorted cell type to the tumors normalized to one. Samples are sorted according to figure 1A **C.** *t*-SNE plot depicting PanNET and normal α- and β-cell samples. In the blue and orange circles are indicated α- and β-cell samples, respectively (2 samples for each cell type). Orange and blue points indicate β-like and α-like tumour, respectively.

**Figure S3.** In fig. **A** and **B** illustrative pictures for Cenp11 (green) and MEN1 (red) FISH. In fig. **C** and **D** and illustrative pictures for Cenp17 (red) and Cenp4 (green) FISH**. E.** Dominant inferred copy number (CN) for each chromosomal arm in each tumour. Chromosome arms in rows and patients in columns (red: amplification, blue: deletions, cut-off at ±0.2). CNA group 1 with limited number of events; group 2 with losses and gains; group 3 with recurrent chromosomal losses.

**Figure S4.** ARX and PDX1 IHC mark respectively α- and β-cells in normal pancreatic islets (drawn in green). Many exocrine and ductal cells present positivity for PDX1, as known[1–4].

**Figure S5. A.** Number of PanNETs (y-axis) per cell-type subtype (x-axis) and relative to tumour stage **B.** Number of PanNETs (y-axis) per cell-type subtype (x-axis) and relative to patient relapse. **C.** Kaplan-Meier disease free survival of 34 patients (cohort 2) stratified according to PDX1, ARX and DAXX/ATRX IHC.

**Figure S6. A.** Plot of cumulative distribution functions (CDF) for the consensus matrix for each *k* (left) and relative change in area under the CDF curve (right). Consensus clustering was performed for the 125 PanNETs (cohort 1) according to the 6364 differentially methylated sites between α-like, β-like and intermediate tumours (adj. p-value<0.001 and $|\Delta\beta|>0.2$). **B.** Kaplan-Meier disease free survival of the 98 patients (cohort 1) stratified according the consensus clustering groups (α-like, β-like, intermediate-WT, intermediate-ADM). **C.** Plot of cumulative distribution functions (CDF) for the consensus matrix for each *k* (left) and relative change in area under the CDF curve (right). Consensus clustering was performed for the 157 PanNETs (cohort 1 and Chan *et al.* cohort) according to the 6359 differentially methylated sites (5 sites were excluded after filtering and normalization processes) identified from the analysis of cohort 1. In **D.** the relative consensus clustering matrix for *k*=4. Each column represents a patient. Consensus cluster correlation is indicated according to the blue scale as indicated. **E.** Kaplan-Meier disease free survival of the 122 patients (cohort 1 and Chan *et al.* cohort) stratified according the consensus cluster groups (α-like, β-like, intermediate-WT, intermediate-ADM).

### Figure S1

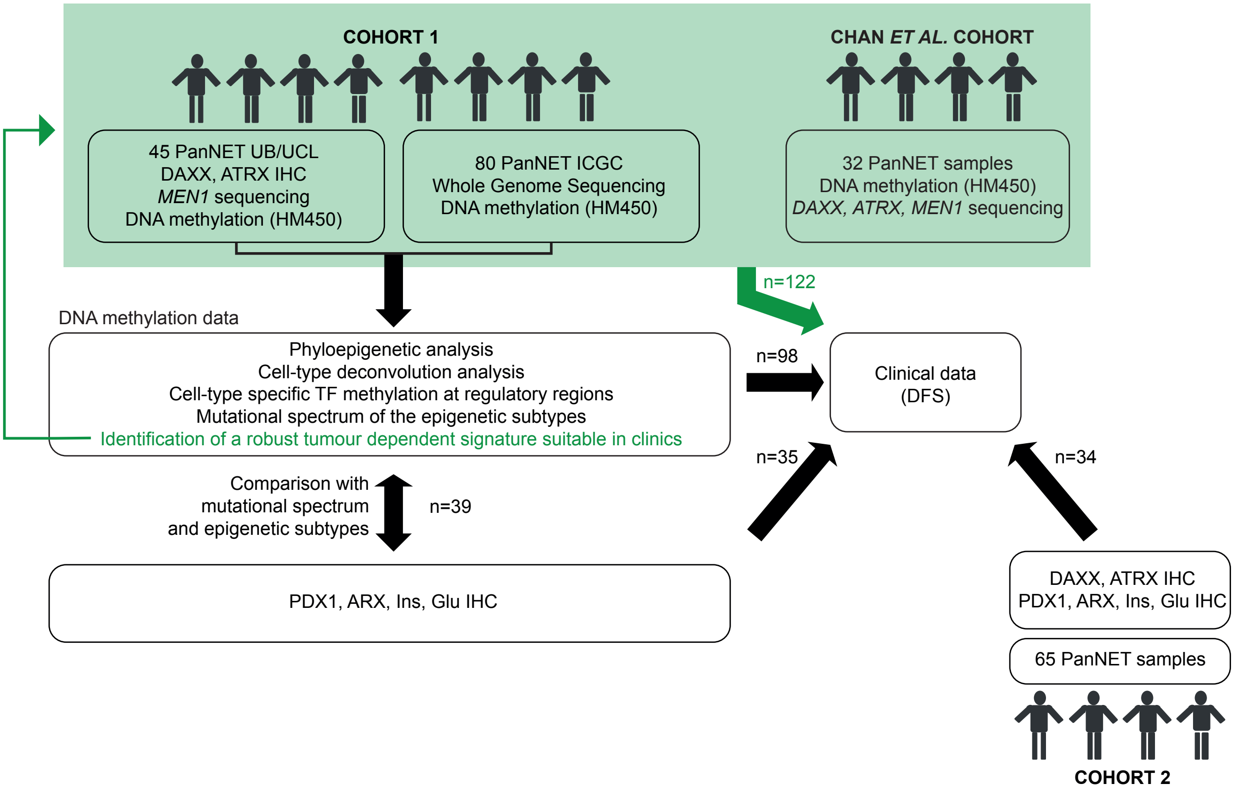
COHORT 1
CHAN ET AL. COHORT
45 PanNET UB/UCL
DAXX, ATRX IHC
MEN1 sequencing
DNA methylation (HM450)
80 PanNET ICGC
Whole Genome Sequencing
DNA methylation (HM450)
32 PanNET samples
DNA methylation (HM450)
DAXX, ATRX, MEN1 sequencing
n=122
DNA methylation data
Phyloepigenetic analysis
Cell-type deconvolution analysis
Cell-type specific TF methylation at regulatory regions
Mutational spectrum of the epigenetic subtypes
Identification of a robust tumour dependent signature suitable in clinics
n=98
Clinical data
(DFS)
n=35
n=34
Comparison with
mutational spectrum
and epigenetic subtypes
n=39
PDX1, ARX, Ins, Glu IHC
DAXX, ATRX IHC
PDX1, ARX, Ins, Glu IHC
65 PanNET samples
COHORT 2

### Figure S2

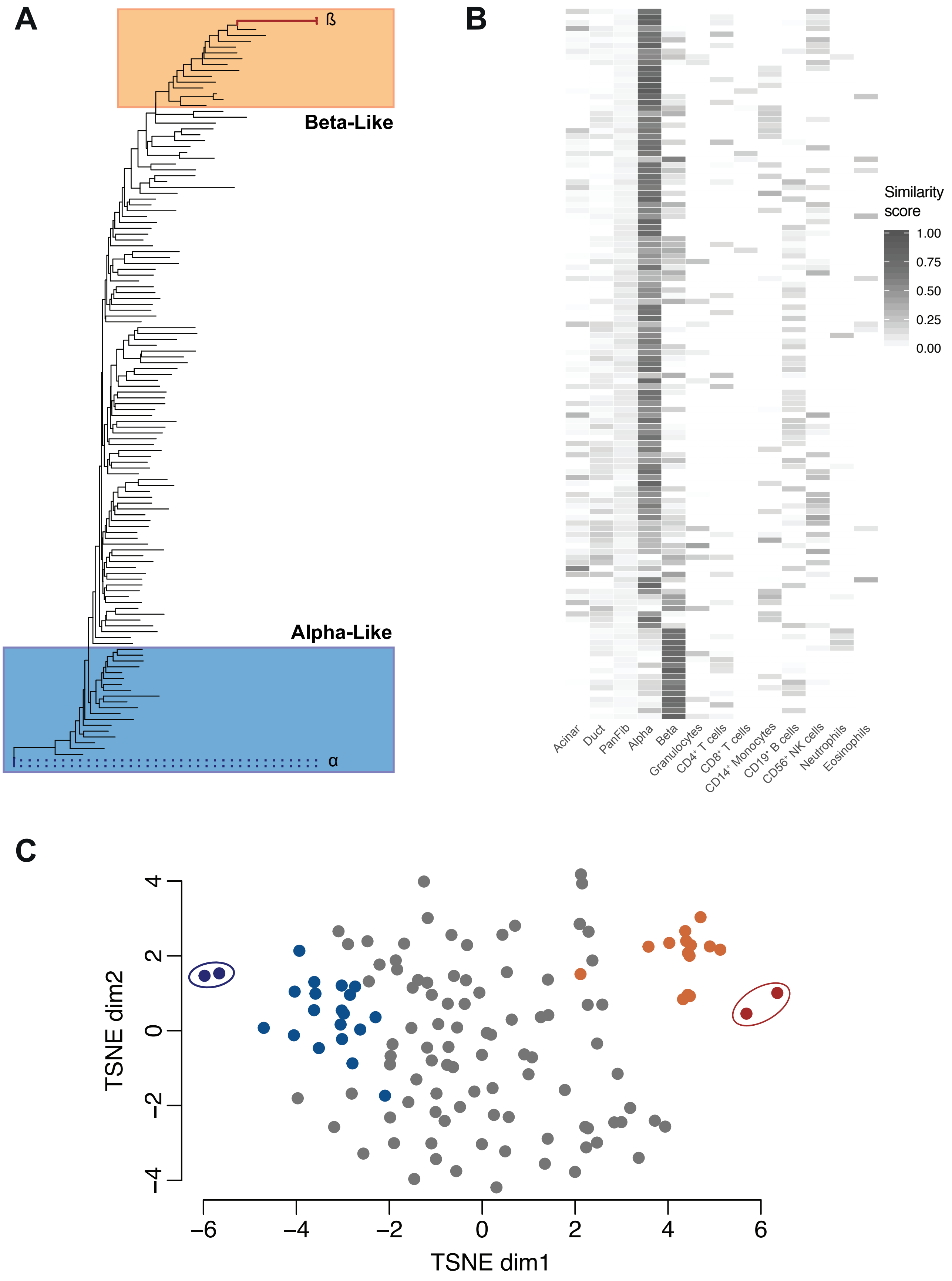

A
ß
Beta-Like
Alpha-Like
α
B
Similarity score
1.00
0.75
0.50
0.25
0.00
Acinar
Duct
PanFib
Alpha
Beta
Granulocytes
CD4+ T cells
CD8+ T cells
CD14+ Monocytes
CD19+ B cells
CD56+ NK cells
Neutrophils
Eosinophils
C
TSNE dim2
TSNE dim1
4
2
0
−2
−4
−6
6

### Figure S3

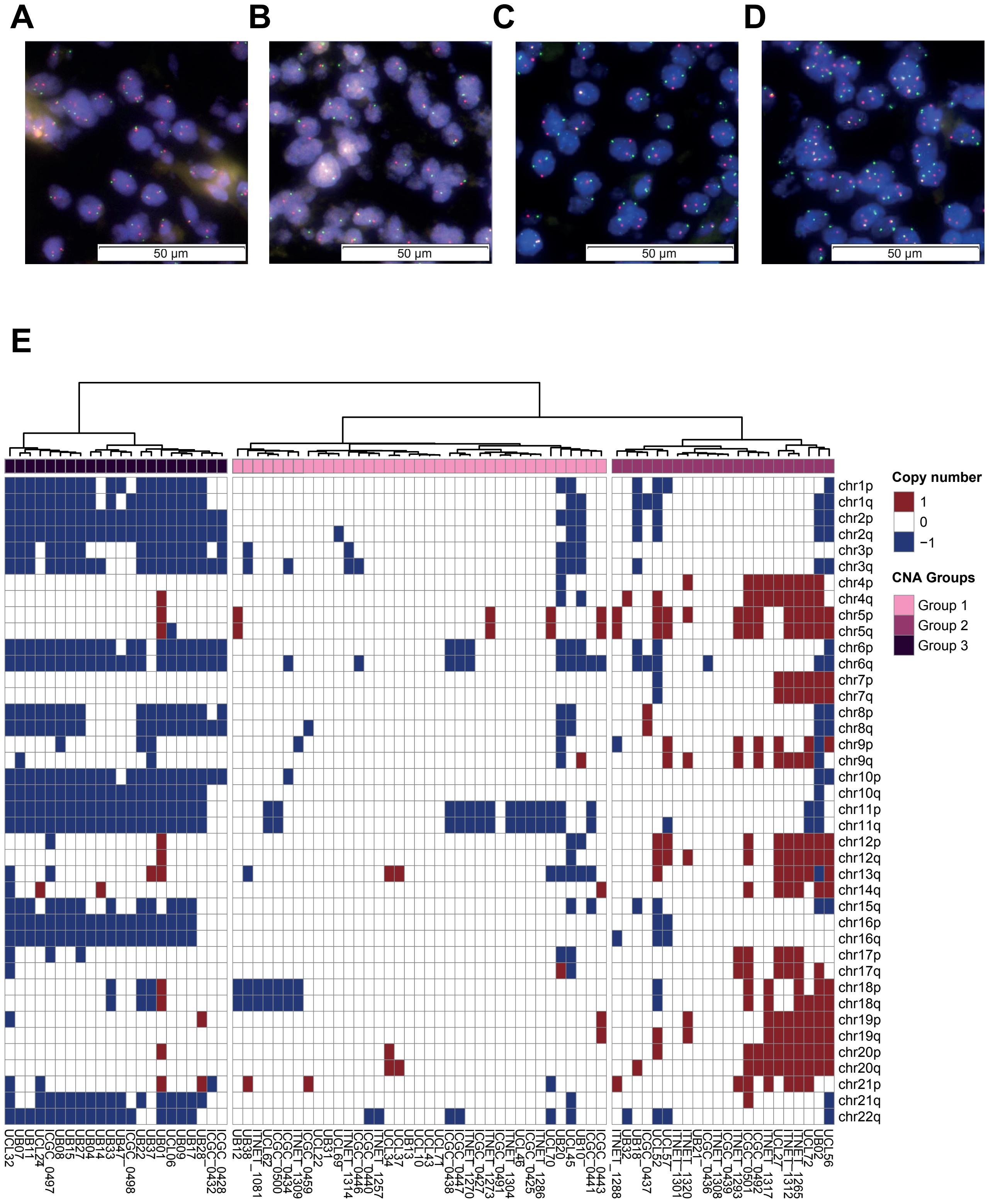

A
B
C
D
50 µm
50 µm
50 µm
50 µm
E
Copy number
1
0
−1
CNA Groups
Group 1
Group 2
Group 3
chr1p
chr1q
chr2p
chr2q
chr3p
chr3q
chr4p
chr4q
chr5p
chr5q
chr6p
chr6q
chr7p
chr7q
chr8p
chr8q
chr9p
chr9q
chr10p
chr10q
chr11p
chr11q
chr12p
chr12q
chr13q
chr14q
chr15q
chr16p
chr16q
chr17p
chr17q
chr18p
chr18q
chr19p
chr19q
chr20p
chr20q
chr21p
chr21q
chr22q

Figure S4

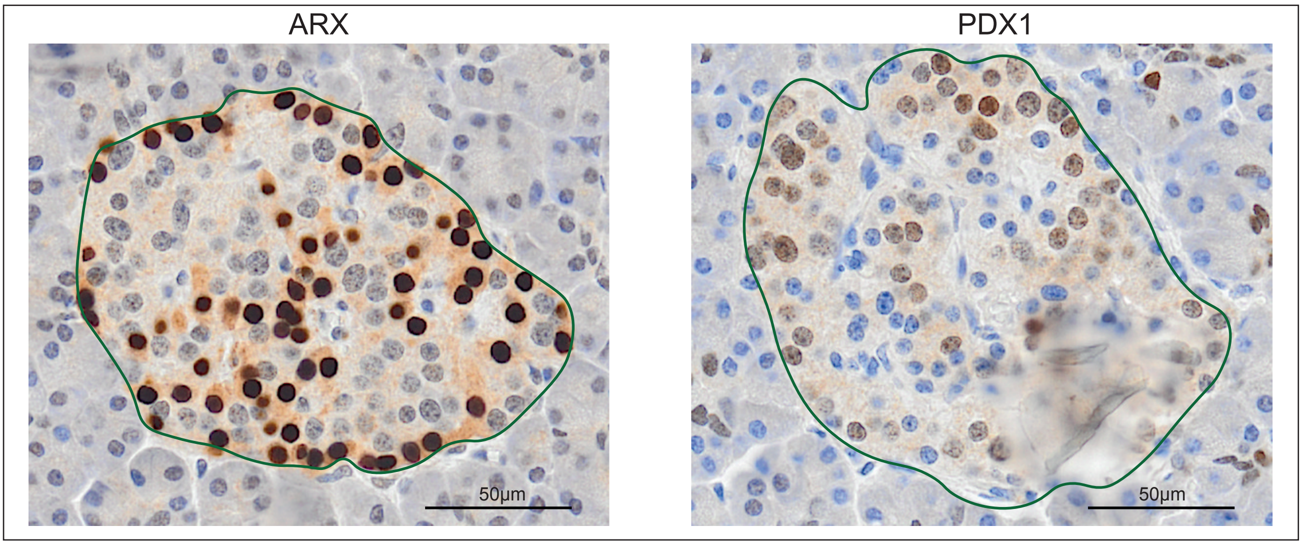

### Figure S5

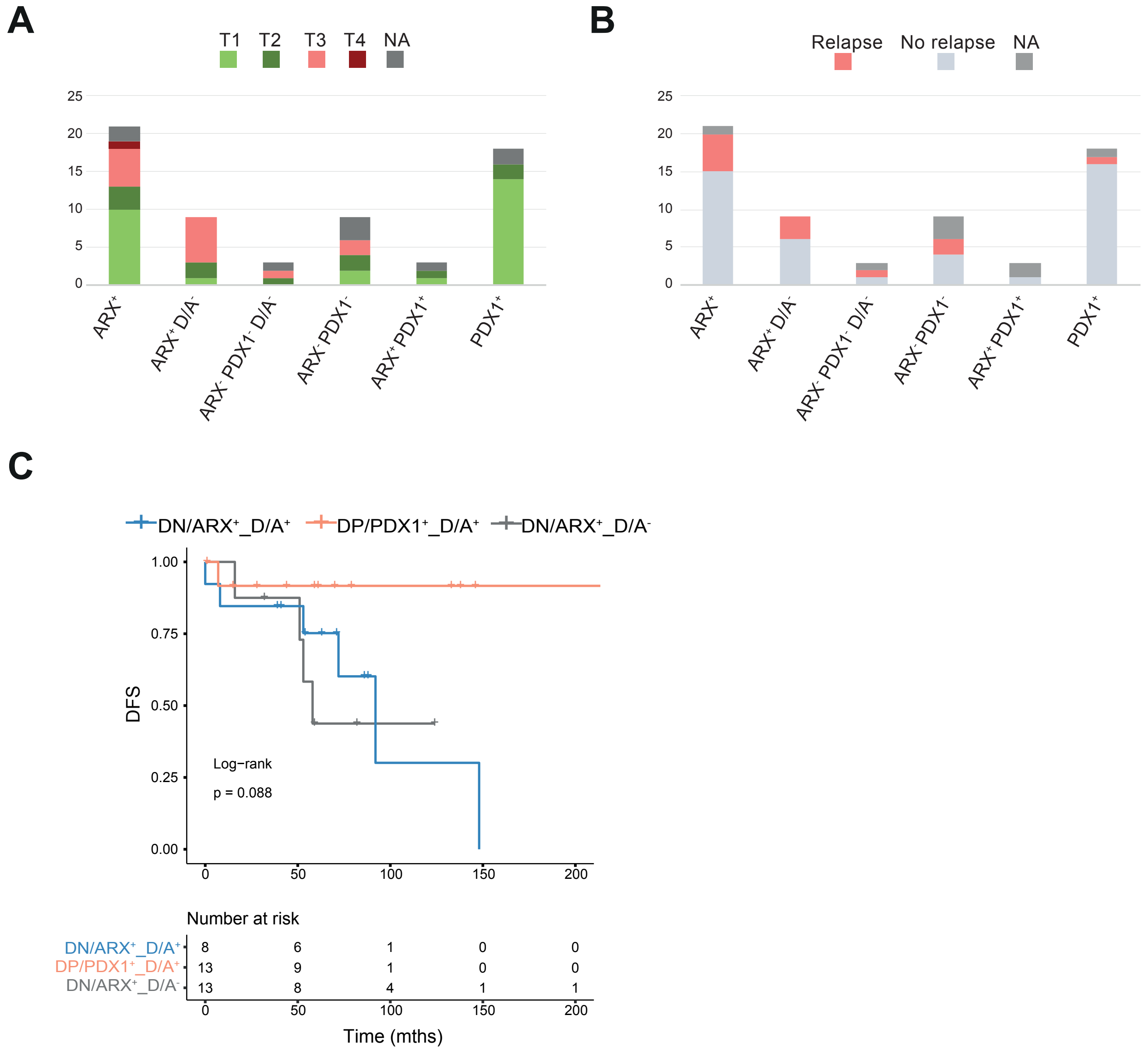
A
T1 T2 T3 T4 NA
25
20
15
10
5
0
ARX+
ARX+D/A-
ARX- PDX1- D/A-
ARX-PDX1-
ARX+PDX1+
PDX1+
B
Relapse No relapse NA
25
20
15
10
5
0
ARX+
ARX+D/A-
ARX- PDX1- D/A-
ARX-PDX1-
ARX+PDX1+
PDX1+
C
DN/ARX+_D/A+ DP/PDX1+_D/A+ DN/ARX+_D/A-
1.00
0.75
0.50
0.25
0.00
DFS
Log−rank
p = 0.088
0 50 100 150 200
Number at risk
DN/ARX+_D/A+ 8 6 1 0 0
DP/PDX1+_D/A+ 13 9 1 0 0
DN/ARX+_D/A- 13 8 4 1 1
0 50 100 150 200
Time (mths)

### Figure S6

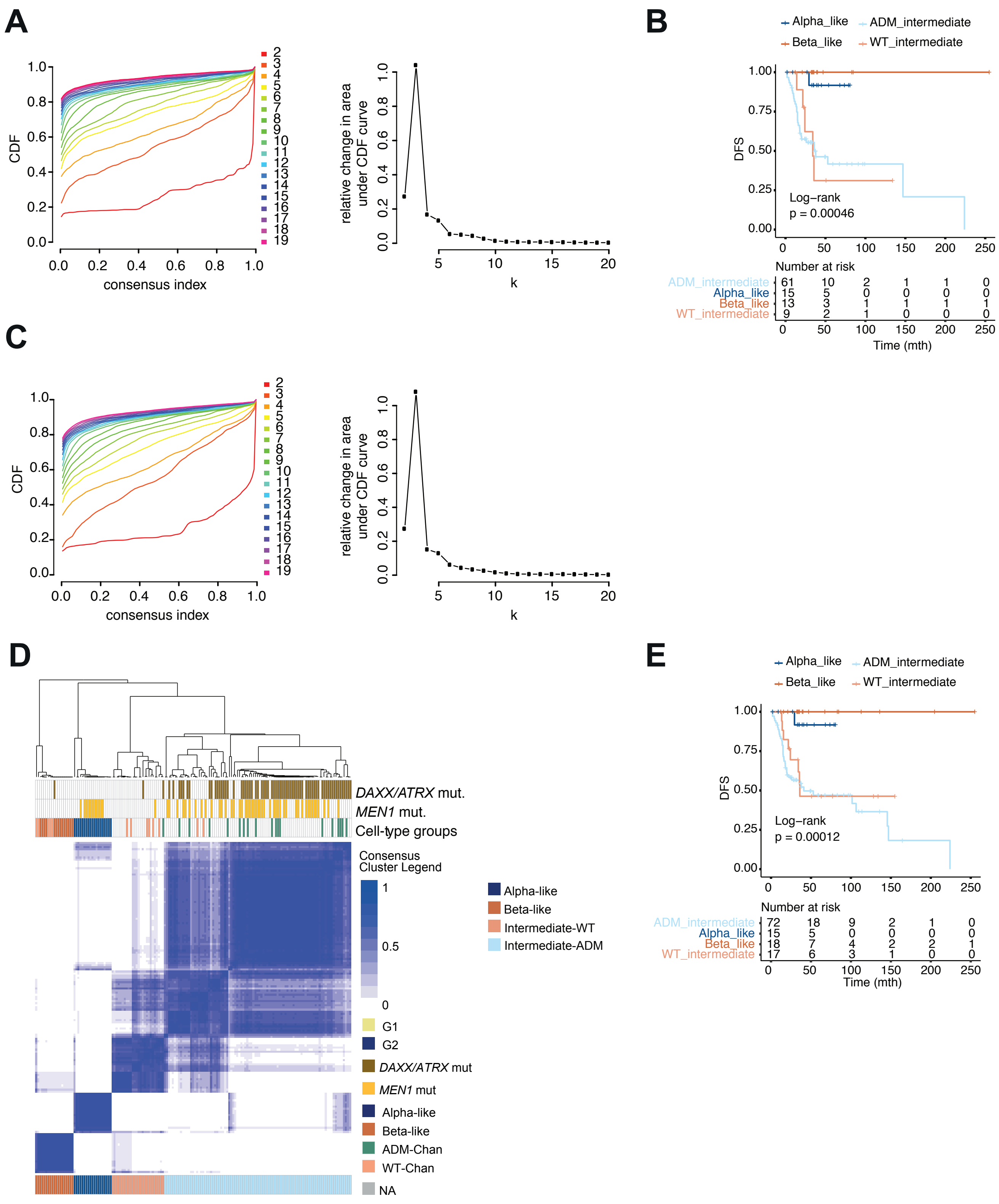
A
B
C
D
E
CDF
consensus index
relative change in area under CDF curve
k
Alpha_like
Beta_like
ADM_intermediate
WT_intermediate
DFS
Log−rank
p = 0.00046
Number at risk
ADM_intermediate 61 10 2 1 1 0
Alpha_like 15 5 0 0 0 0
Beta_like 13 3 1 1 1 1
WT_intermediate 9 2 1 0 0 0
Time (mth)
DAXX/ATRX mut.
MEN1 mut.
Cell-type groups
Consensus Cluster Legend
1
0.5
0
G1
G2
DAXX/ATRX mut
MEN1 mut
Alpha-like
Beta-like
ADM-Chan
WT-Chan
NA
Alpha-like
Beta-like
Intermediate-WT
Intermediate-ADM
Log−rank
p = 0.00012
Number at risk
ADM_intermediate 72 18 9 2 1 0
Alpha_like 15 5 0 0 0 0
Beta_like 18 7 4 2 2 1
WT_intermediate 17 6 3 1 0 0
Time (mth)
